# “Evaluating the Benefits and Limits of Multiple Displacement Amplification with Whole-Genome Oxford Nanopore Sequencing”

**DOI:** 10.1101/2024.02.09.579537

**Authors:** Fiifi A. Dadzie, Megan S. Beaudry, Alex Deyanov, Haley Slanis, Minh Q. Duong, Randi Turner, Asis Khan, Cesar A. Arias, Jessica C. Kissinger, Travis C. Glenn, Rodrigo de Paula Baptista

## Abstract

Multiple Displacement Amplification (MDA) outperforms conventional PCR in long fragment and whole genome amplification which makes it attractive to couple with long-read sequencing of samples with limited quantities of DNA to obtain improved genome assemblies. Here, we explore the efficacy and limits of MDA for genome sequence assembly using Oxford Nanopore Technologies (ONT) rapid library preparations and minION sequencing. We successfully generated almost complete genome sequences for all organisms examined, including *Cryptosporidium meleagridis*, *Staphylococcus aureus*, *Enterococcus faecium*, and *Escherichia coli,* with the ability to generate high-quality data from samples starting with only 0.025 ng of total DNA. Controlled sheared DNA samples exhibited a distinct pattern of size-increase after MDA, which may be associated with the amplification of long, low-abundance fragments present in the assay, as well as generating concatemeric sequences during amplification. To address concatemers, we developed a computational pipeline (CADECT: Concatemer Detection Tool) to identify and remove putative concatemeric sequences. This study highlights the efficacy of MDA in generating high-quality genome assemblies from limited amounts of input DNA. Also, the CADECT pipeline effectively mitigated the impact of concatemeric sequences, enabling the assembly of contiguous sequences even in cases where the input genomic DNA was degraded. These results have significant implications for the study of organisms that are challenging to culture *in vitro*, such as *Cryptosporidium*, and for expediting critical results in clinical settings with limited quantities of available genomic DNA.

## INTRODUCTION

The advent of next-generation sequencing technologies has revolutionized genomics research by enabling the rapid and cost-effective generation of vast amounts of sequencing data (Slatko et al. 2018; Hu et al. 2021). Among these technologies, Oxford Nanopore Sequencing (ONT) stands out due to its ability to provide long-read sequencing data in real-time, with lower instrument costs and less input DNA required for non-amplified library preparations than the other major commercial long-read sequencing platform, PacBio (Pacbio 2022). ONT sequencing has been used for numerous applications, including *de novo* genome assembly, metagenomics, and pathogen detection. However, ONT sequencing library preparations typically still requires higher-quality and higher quantities of DNA inputs than may be available for many projects. ONT rapid library preparations usually require at least 50 ng input per sample, but more is required when pooling with <8 other barcoded samples (≥400 ng is recommended for loading onto a MinION flow cell). Many samples also suffer from DNA degradation, where the majority of DNA fragments are shorter than is desirable for ONT library preparation and removal of small fragments further reduces the quantity of DNA available. This poses challenges when working with samples that have limited quantity and/or degraded DNA (Delahaye and Nicolas 2021). For this reason, alternative library preparation or sequencing techniques, including short-read sequencers (*e.g*., Illumina, Element Biosciences AVITI), are often preferred for handling samples with low molecular weight and/or low quantities of DNA.

To overcome these limitations, multiple displacement amplification (MDA) has emerged as a valuable and highly efficient method for amplifying small quantities of DNA. MDA has significant advantages over conventional PCR and other whole genome amplification techniques (Hou et al. 2015). These advantages include reduced waste of rare samples, isothermal amplification for efficiency, heightened sensitivity in detecting low amounts of DNA inputs, minimized bias and error rates, amplification of long DNA fragments and whole genome amplification of organisms with relatively small genome size (< 10Mb). MDA utilizes the Phi29 DNA polymerase with a displacement activity that enables the isothermal amplification of DNA with high fidelity and exponential amplification of DNA molecules (Dean et al. 2002). This technique has been successfully applied in various genomic studies, including single-cell sequencing, ancient DNA analysis, and microbiome studies (Binga et al. 2008; Lasken 2009). Moreover, MDA enables the amplification of long DNA fragments, making it valuable for applications such as cloning and genomic library preparation (Fullwood et al. 2008). While a protocol for MDA with ligation sequencing kits (Qiagen, Germany) is available, MDA’s application with ONT rapid kits, which offer faster processing times and yield relatively smaller fragments compared to ligation kits, has not been extensively investigated. Consequently, MDA’s potential limitations and impacts on whole-genome assembly in this context remain relatively unexplored.

The use of MDA combined with ONT sequencing has the potential to unlock genomic insights for organisms that are small (e.g., larval ticks, parasitoid wasps, etc.) to microscopic, especially those that are difficult or impossible to culture *in vitro* (*e.g*. *Cryptosporidium* species, *Mycobacterium leprae* and *Treponema pallidum*). Furthermore, clinical samples and isolates with limiting amounts of DNA pose a challenge for rapid and accurate genome sequence analysis, especially in urgent clinical situations where timely results are crucial. Working with degraded DNA samples becomes an issue since it could limit the sequence genomic coverage and assembly (Ceccherini et al. 2003). MDA is not suitable for analysis of severely degraded DNA, since could impact: (i) MDA efficiency due potential breaks or lesions leading incomplete or suboptimal amplification; (ii) bias resulting in uneven coverage across the genome; and (iii) contaminants that could interfere with the MDA reaction (Wang et al. 2004).

It’s important to mention that when utilizing MDA there are limitations that needs to be considered to ensure the reliability and integrity of the sequencing results. While MDA has facilitated genomic sequencing from low concentrations of template nucleic acid, there are still several limitations to consider. These include: (i) Nonspecific amplification resulted from primer dimer formation causing template switching or contamination by DNA templates; (ii) Formation of chimeric DNA rearrangements; and (iii) Representation bias, which can affect the accuracy and completeness of the amplified genomic material (Binga et al. 2008). Some studies shows that chimeric reads are usually invert chimeras or direct chimeras, but it was previously observed that most of detected MDA chimeric sequences (85%) are inverted chimeras, such as inverted sequences with intervening deletions which can be caused by template switching (Lasken and Stockwell 2007; Lu et al. 2023). These chimeric sequences are known to affect genome sequencing since they can be considered as amplification artifacts, which cannot be used for genome assembling (Lu et al. 2023). Studies suggest that chimerism in MDA sequencing data is a significant concern that is gaining attention, particularly with the rise of single-cell studies (Hard et al. 2023).

To address the challenges associated with artifactual concatemeric sequences generated during MDA, we developed a novel bioinformatic tool called CADECT (Concatemer Detection Tool), which is made available at https://github.com/rpbap/CADECT. This tool enabled the identification and removal of putative inverted chimeric concatemers, thus improving the accuracy and contiguity of the genome assembly.

Our study aims to provide valuable insights into the use of MDA for whole-genome ONT sequencing, particularly for low molecular weight and/or low quantities of DNA samples, highlighting its potential as a powerful method to obtain high-quality long-read sequencing data. We assessed the MDA advantages and constraints, and effectiveness for whole-genome assembly in microbial organisms with genome sizes <10 Mb. This is especially significant for infectious disease agents, where obtaining enough DNA can be challenging. Overall, our study underscores the potential of MDA in enabling high-quality long-read sequencing from challenging low-concentration DNA samples, emphasizing its importance in various genomic research and clinical applications.

## RESULTS

### WGA results

Our whole genome amplification (WGA) results reveal that in each sample type tested, we find an overall fold change of > 500× in comparison to the original sample (Table 1). Following amplification, approximately 1.5 µg of the product was debranched using T7 endonuclease prior to library preparation for ONT sequencing. Typically, we experience a ∼45% recovery after this step, attributed to the bead purification process (Table 1). Though a significant amount of DNA is lost during the DNA purification step post T7 endonuclease reaction, an overall fold change of ∼100× is observed when compared to the WGA DNA input.

**Table 1.** Observed amplification yield increase by sample type.

|  | WGA input<br>(ng) | WGA output<br>(ng) | T7 output<br>(ng) | T7 recovery<br>(%) | Estimated<br>Fold<br>Increase |
| --- | --- | --- | --- | --- | --- |
| Gram-positive<br>(*S. aureus*) | 2.5 | 1500 | 894 | 59.6 | 357.6× |
| Gram-negative (*E. coli*) | 5.0 | 8360 | 552 | 36.8 | 110.0× |
| Eukaryotic<br>Pathogen<br>(*Cryptosporidium* ssp.) | 2.5 | 1976 | 555 | 44.1 | 222.0× |
| Background<br>(Calf thymus) | 5.0 | 3800 | 620 | 41.33 | 124.0× |

We successfully obtained contiguous and sometimes even chromosomal-level assemblies from the samples analyzed in this study, starting with DNA inputs significantly lower than Oxford Nanopore’s recommended minimum of 50 ng for the rapid barcode kit (Table S1).

For certain samples, such as *E. faecium*, we observed that achieving improved contiguity required generating higher depth coverage during the sequencing. Our results indicate that, for this organism, reaching depths beyond 70× allowed us to attain a chromosomal-level assembly with only 2.5 ng of starting total DNA (Table S2). In comparison, increased sequencing depth on samples that started with less than 0.001 ng of input into the MDA did not enhance contiguity. Combining separate MDA amplifications of the same limited input samples did improve the final genome coverage because the random nature of the initial templates and amplification process. Thus, multiple independent MDAs appears to be advantageous because it could randomly amplify by chance different regions that are beneficial for the genome assembly.

To check for potential GC bias on the sequencing depth along the genome, the R^2^ correlation coefficient between average depth and average %GC across 1000 base pair regions of the *E. faecium* non-amplified and amplified assemblies was 0.0262 and 0.0265, respectively (Fig. S1).

### WGA amplification using serially diluted samples

Serial dilutions of a single *E. faecium* sample reveal successful DNA amplification even at low initial DNA amount of 2.5E-5 ng (Fig. 1A). The MDA technique imposes a size limit on its amplified products, with an average product length of 10-12 kb (Dean et al. 2002), and it requires a debranching step, leading to a reduction in the mean sequence read sizes (Table S3). Post MDA, the average size of the reads typically falls within the range of 2-3 kb. In contrast, standard Oxford Nanopore Technologies (ONT) assays without amplification (*i.e*., ONT Rapid Barcode Kit (RBK)) which includes a transposase step that simultaneously cleaves template molecules and attaches tags to the cleaved ends, typically generate DNA fragments ranging from 5-20 kb. However, when assessing genome coverage (genome sizes <10 Mb), we observed that DNA inputs below 0.025 ng result in incomplete coverage of certain genomic regions (Fig. 1B).

**Figure 1.**
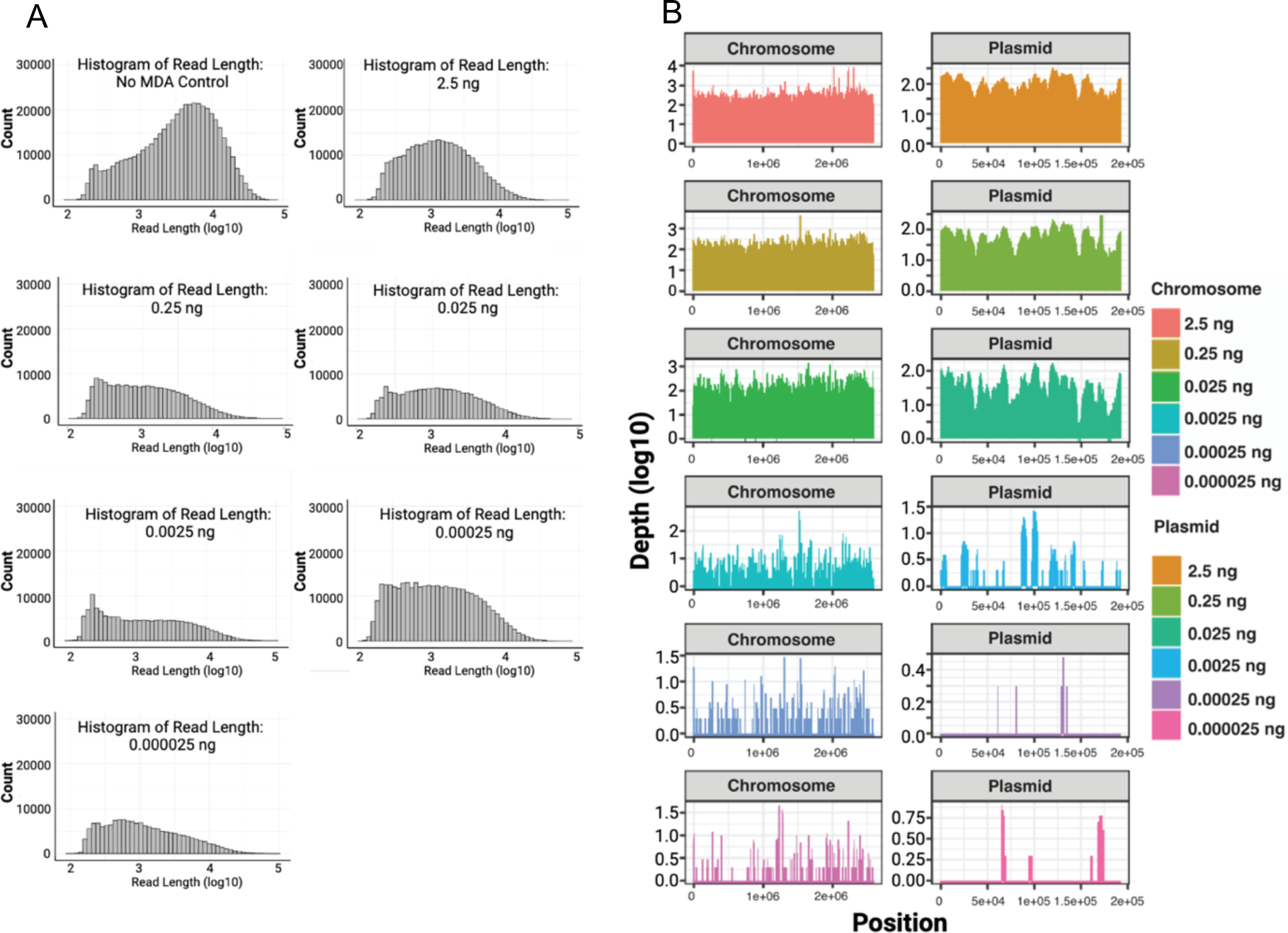
Serial dilution test with *E. faecium* sample DNA. (A) Depicts the distribution and counts (y-axis) of read lengths (x-axis) across all diluted samples. (B) Illustrates the horizontal coverage of the chromosomal regions across all diluted and subsequently amplified samples, with read depth on the y-axis and genome position on the x-axis.

### MDA results in an unexpected size increase from fragmented DNA samples

Following controlled enzymatic fragmentation using a dsDNA fragmentase and MDA according to our protocol, we observed an unexpected size increase distribution of fragments (Fig. 2A). Indeed, for all samples, except *Cryptosporidium*, the size distribution post-MDA was nearly identical for intact and fragmented input DNA. Subsequent analysis of the ONT sequencing results revealed the existence of read lengths that are longer than those present in the same sample without MDA amplification (Fig. 2B).

**Figure 2.**
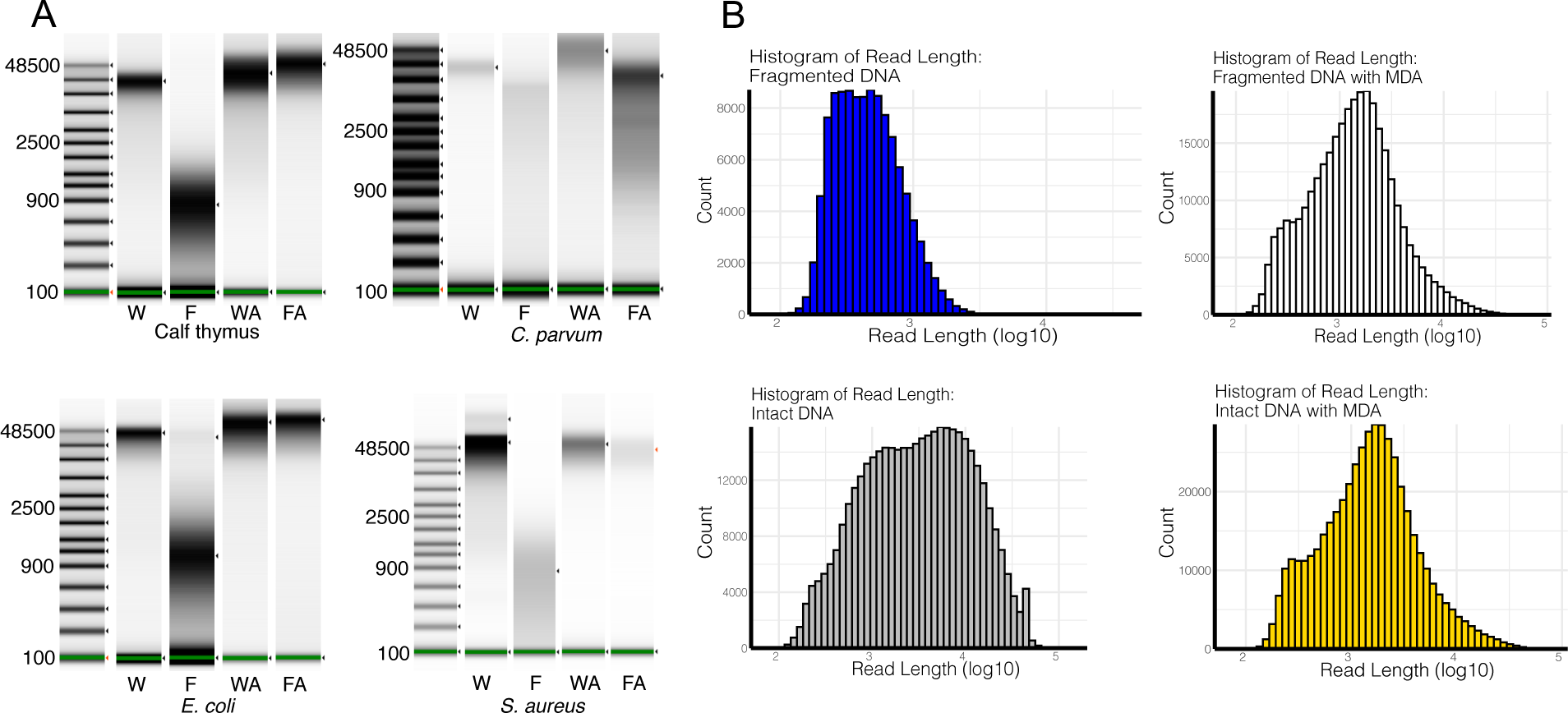
DNA fragment and read Size Range pre- and post-Multiple Displacement Amplification using Size-Controlled Fragmented DNA. (A) TapeStation results for different organism sets; (B) ONT sequencing results obtained before and after amplification for *S. aureus*. W = whole intact DNA; F = Fragmented DNA; WA and FA = after amplification; and WT and FT = after T7 debranching. Uncropped TapeStation results are in Fig. S2.

Upon closer examination of the sequence content, two distinct types of reads were identified. Some represented potentially low-abundance longer reads that escaped fragmentation during the enzyme incubation and were subsequently amplified. The other reads were primarily chimeric concatemers, likely generated through template switching of short fragments during MDA (Fig. 3A). While the occurrence of concatemers in MDA assays has been reported previously (Paul and Apgar 2005; Lu et al. 2023), they are typically present in low amounts after sequencing. In our case, the fragmentation process seemed to enhance the prevalence of these chimeric reads in our ONT sequencing. As expected, assembly of the data revealed that the presence of these chimeric/concatemers regions significantly impacted genome assembly, resulting in bubble fragmentation effect across the entire genome and affecting contiguity (Fig. 3B).

**Figure 3.**
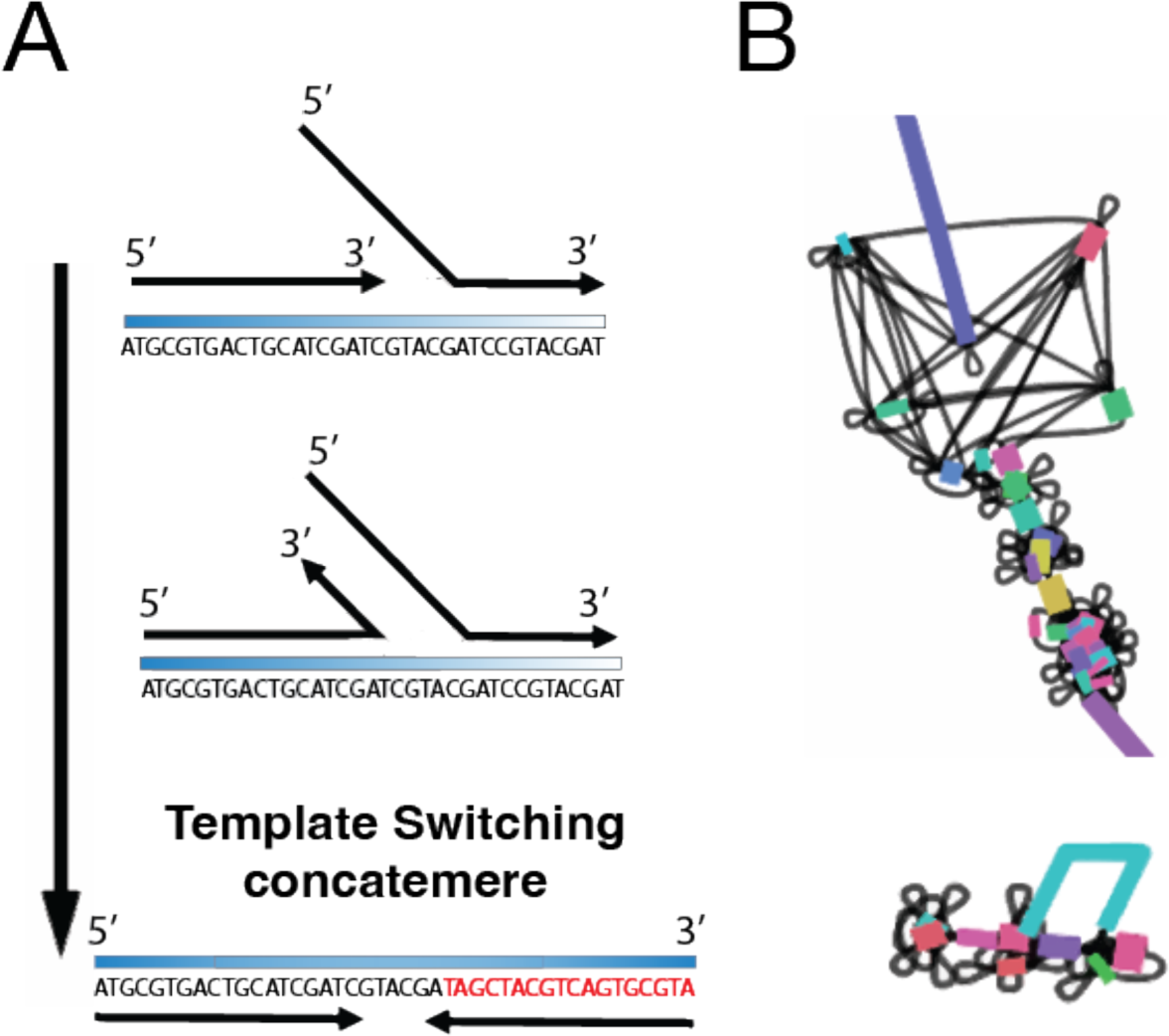
Impact of MDA-Generated Concatemers on the Genome Assembly. (A) Concatemers generated by template switching; (B) Graph representation of the effect of concatemers on genome assembly (bubble fragmentation effect).

### Concatemer detection tool

To identify and eliminate potential concatemers generated by MDA, we designed a concatemer detection tool specifically tailored for raw ONT reads called CADECT. This tool enables the differentiation of putative concatemeric chimeric reads from non-concatemeric ones. To achieve this, the process involves dividing each long-read sequence into multiple fragments using a sliding window approach and then aligning these fragments with one another. The underlying hypothesis is that the presence of a concatemer would result in certain windows aligning with each other, thereby confirming the existence of a potential concatemer or tandem repeat within the sequenced read. Reads with lengths less than twice the given window size are categorized and stored as short reads. Additionally, it incorporates a size selection mechanism to isolate longer reads, thereby streamlining the genome assembly process and enhancing contiguity.

Following evaluation of the CADECT pipeline on fragmented DNA and a comparative analysis of results pre- and post-amplification assay (Table S1), we confirmed that the final genome assembly exhibited significantly reduced fragmentation. The integration of MDA and CADECT proved to be effective, particularly in handling challenging, low quantity DNA samples. This combination facilitated the generation of nearly complete genome assemblies with depths above 70× (Fig. 4; Table S4).

**Figure 4.**
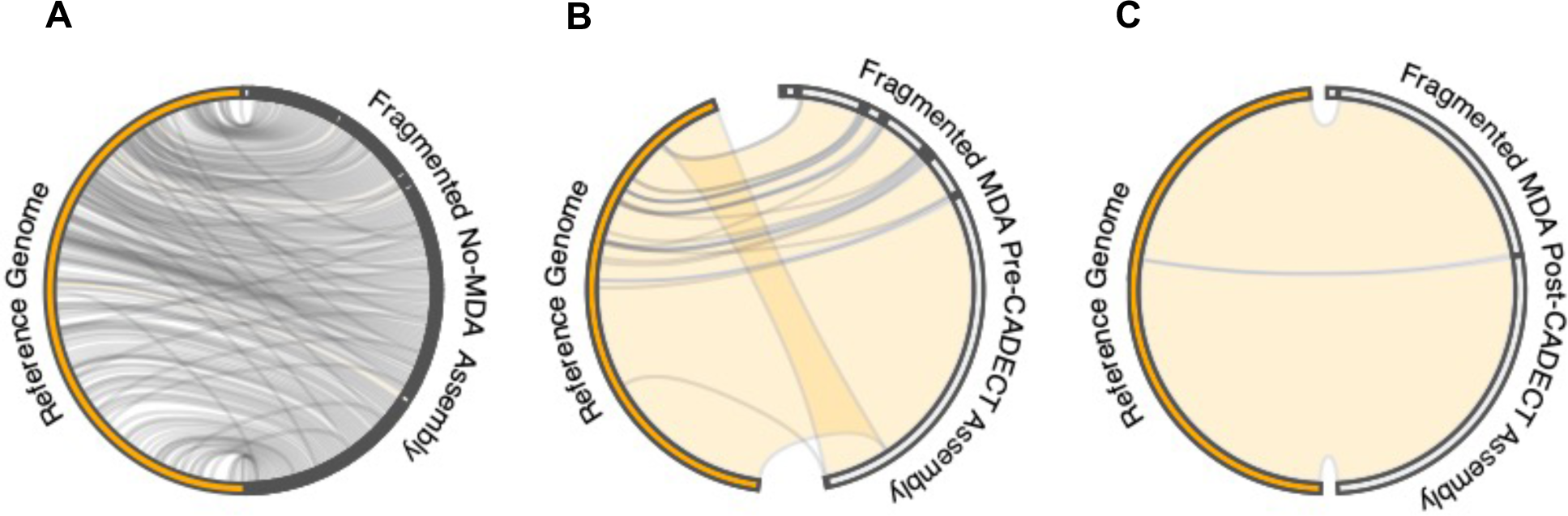
Circos plot illustrating a synteny comparison between the reference *S. aureus* ATCC-29213 genome sequence and pre- and post-amplification genome assemblies. The Circos plots contrast the assemblies resulting from the amplified fragmented DNA before and after CADECT processing. A comparison between the reference genome and (A) genomic assembly of fragmented DNA sample without MDA, (B) genomic assembly of fragmented DNA sample with MDA before CADECT and (C) genomic assembly of fragmented DNA sample with MDA after CADECT.

Overall, when comparing the data before and after CADECT using default parameters with a 500 base window size, we observe that its stringent process, which separates putative concatemers and shorter reads, tends to affect the average final depth of the final input. Specifically, in the case of *S. aureus*, we note that for high-quality intact amplified DNA, the detection of putative chimeras and size selection decreases coverage by 40%, whereas for amplified fragmented samples, it decreases coverage by 50% (Table S5). The effect on depth is more pronounced for fragmented samples due to size selection.

## DISCUSSION

Our study demonstrates that MDA offers a promising solution for amplifying low amounts of DNA of precious samples for ONT sequence generation with the ONT rapid barcode sequencing Library kit (RBK). In this study, we demonstrated three key points: (A) Using our method, we can successfully sequence samples with DNA inputs as low as 0.025 pg. This suggests that long-read sequencing of single cells may be possible. Single-cell sequencing represents a significant achievement as variability, if it exists in the sample, is reduced in the sequence results because we are targeting a considerably smaller number of cells compared to larger bulk-extracted samples. (B) For *Cryptosporidium*, we show that we can reach single-oocyst levels, as 0.025 pg is equivalent to ∼1.5 times the amount of DNA in one oocyst (Table 2). This is significant for *Cryptosporidium* because this parasite cannot be cloned. The prospect of Single-oocyst sequencing removes the variation introduced with bulk sequencing approaches. (C) We explored and showed the potential of Whole Genome Amplification (WGA) as an option to examine even smaller quantities of DNA depending on the organism under investigation and the size of its genome. Here, we have only examined organisms with genome sizes < 10Mb. Larger genome sizes will require additional starting material and smaller genome sizes should have success with even less input DNA.

**Table 2.** Estimated DNA concentration in a single cell of the organisms studied in this project.

| SAMPLE | Estimated genome size (Mb) | Amt of DNA/cell (fg) |
| --- | --- | --- |
| *E. coli* | 5.00 | 5.43 |
| *S. aureus* | 2.81 | 3.03 |
| *E. faecium* | 2.91 | 3.14 |
| *Cryptosporidium* ssp. (oocysts)* | 9.2 | 39.70 |
| *Cryptosporidium* ssp. (sporozoite) |  | 9.90 |

\*One *Cryptosporidium* ssp. oocyst contains 4 haploid sporozoites.

We were able to obtain whole genome sequences at the chromosomal level for almost all tested organisms when generating depth coverages >70×. This indicates that we can overcome the challenge of the relatively shorter reads that MDA with T7 debranching generates, in comparison to reads generated from higher DNA input without amplification. It’s essential to highlight that highly complex regions, such as repetitive regions with tandem repeats larger than the window size used for CADECT detection, might be identified as concatemeric reads. This occurs because the tool detects repeat overlaps, which can lead to their exclusion before the assembly process, potentially causing some regions of the genome to remain fragmented. This outcome may vary depending on the organism being sequenced.

At a lower amount of initial DNA, we observed that the amplification appears to be random, exhibiting no apparent bias across the genome (Fig. 1). At extremely low DNA amounts, achieving full coverage of the target genome may be challenging, but the method remains valuable for potential taxon identification and may prove effective for the identification of plasmids as well (Fig. 1). While the effectiveness in metagenomic samples requires further evaluation, there is promise in using this approach for taxon identification. Extremely low-abundance samples tend to produce patchy sequence information. Thus, although extremely low-concentration samples provide valuable sequence information, they also lack coverage in many regions, which impacts the assembly process and the ability to produce full genomic information. Combining multiple MDA replicates is likely to increase the chances of amplifying more regions and thus will be more likely to enhance genome coverage verses deeper sequencing. Interestingly, GC ratios apparently did not impact the amplification showing very low correlations (Fig. S1).

In the case of sheared DNA, the higher impact on the depth after CADECT is primarily related to loss in the size selection pipeline (Table S5). However, our concatemer detection tool, CADECT, effectively identified and removed several concatemers, facilitating the assembly and yielding good results. This highlights the importance of bioinformatic tools in overcoming challenges associated with amplification artifacts thus improving the accuracy of genome assembly.

It is worth noting that the CADECT pipeline will remove a significant number of reads which will, impact depth for an optimal genome assembly. If there isn’t sufficient coverage obtained post-CADECT run, an alternative is to merge the reads identified as short by the program with the non-concatemeric reads. As observed previously, the chimeric rate produced by MDA is positively associated with the mean read length (Lu et al. 2023), indicating a decreased likelihood of chimeric reads in this short dataset. Consequently, this dataset is less likely to negatively impact the assembly process. In more complex genome sequences that are rich in repeats, further investigation is required to address these regions effectively and be able to distinguish concatemers from genuine repetitive patterns within the genome. As a solution, the CADECT pipeline generates a separate concatemer fastq file. This file includes putative concatemeric regions as well as true repeats. For example, highly repetitive genomes such as trypanosomatids with an ∼50% genomic repeat content (El-Sayed et al. 2005), CADECT would detect a good number of reads containing real tandem repeats in the genome as putative concatemers, which would result in a higher impact on coverage depth loss and also impact the genome content of the organisms used for assembly. To mitigate this, we recommend incorporating a repeat identification step into the pipeline, such as using RepeatModeler (Flynn et al. 2020) trained with the organism of interest on the putative concatemer generated sequence file from CADECT. This additional step would enhance the recovery of information and data for the subsequent assembly process.

Moving forward, it is crucial to continue exploring the potential of MDA in various biological contexts and optimize the amplification protocol to minimize biases and errors. Additionally, considering the clinical applications of MDA, further research and development of rapid and reliable sequencing approaches are necessary to unlock its full potential in diagnosing and monitoring infectious diseases and other clinical applications.

## MATERIALS AND METHODS

### Sample Collection and Preparation

A 100 ng DNA sample of *Cryptosporidium meleagridis* isolate TU1867 was obtained from BEI Resources (NR-2521) (Manassas, VA). DNA samples from cultured *Staphylococcus aureus* ATCC-29213, *Enterococcus faecium* TX-1330, and *Escherichia coli* strain K12, which were available in our laboratory, were used for testing. The bacterial DNA samples were prepared for downstream processing using the QIAGEN QIAamp DNA Mini Kit with lysozyme for Gram-positive samples and buffer ATL (tissue lysis buffer) for Gram-negatives. To assess sequence integrity, an *S. aureus* DNA sample aliquot was sheared using NEBNext dsDNA Fragmentase for 15 minutes to generate fragments approximately 1000 bp in size. All DNA samples were quality controlled using a TapeStation (Agilent Technologies, Santa Clara, CA) and Qubit (Thermofisher Scientific, Walthma, MA). In addition, we conducted serial dilutions on all samples to assess the limit of detection for amplification in the assay. The dilutions ranged from 2.5E-5 ng to 2.5 ng, allowing us to determine the minimum concentration at which successful amplification could be achieved.

### Multiple Displacement Amplification (MDA)

Prior to whole genome amplification (WGA), the concentration of DNA was obtained using a Qubit fluorometer dsDNA high-sensitivity assay kit (ThermoFisher, Waltham, MA). For the *C. meleagridis* DNA, three different amounts were used as input for whole genome amplification *(i.e*., 2 ng, 5 ng, and 10 ng) in a final volume of 5 µL. For the bacterial samples, MDA was performed on 2.5 ng of fragmented of intact *S. aureus DNA* as well as serial dilutions of DNA from *E. faecium* ranging from 2.5 ng to 2.5E-5 ng. 400 ng of non-amplified DNA was used as an input control for the ONT rapid kit library preparation (Oxford Nanopore Technologies, Oxford, United Kingdom).

Whole genome amplification (WGA) was performed using the Qiagen Repli-G kit (CAT #150023, Qiagen, Hilden, Germany), following the manufacturer’s instructions. Following this, concentrations of DNA were obtained using a Qubit fluorometer dsDNA high-sensitivity assay kit (ThermoFisher, Waltham, MA).

For T7 Endonuclease I debranching, up to 42 µL (*i.e*., all product from the WGA reaction) of WGA DNA was used as input (Catalog #M0302, New England BioLabs, Ipswich, MA). After scaling up the reaction to accommodate a 42 µL input, all reaction components were added following the manufacturer’s guidelines and incubated for 15 minutes at 37°C in a BioRad T100 thermocycler (BioRad, Hercules, CA). The incubated reaction was brought up to a final volume of 50 µL using TE buffer pH 8. AMPure XP beads (CAT# A63880) were prepared ahead of time following the manufacturer’s instructions, and 35 µL of beads were added to the reaction and mixed thoroughly. The bead-reaction mixture was placed on a rotator mixer *(e.g*., Hula mixer) for 10 minutes at room temperature. Following this, the bead-reaction mixture was spun down and placed on a magnet until the eluate was clear and colorless. With the bead-reaction mixture on the magnet, the clear supernatant was pipetted off and 200 µL of freshly prepared 70% ethanol was carefully added not to disturb the pellet (i.e., wash step). The wash step was repeated one time, for a total of two washes. After removing the supernatant from the second wash, 49 µL of water was used to resuspend the pellet which was immediately incubated for one minute at 50°C in a BioRad T100 thermocycler followed by five minutes at room temperature. The bead-reaction mixture was placed back on the magnet, and 49 µL of the elute was transferred to a sterile 1.5 ml tube. Concentrations of DNA were obtained using a Qubit fluorometer dsDNA high-sensitivity assay kit (ThermoFisher, Waltham, MA).

### Whole Genome Sequencing and Assembly

Sequencing of the amplified DNA samples was performed using the ONT SQK-RBK110.96 kit for library preparation R9.4 MinION flow cells (Oxford Nanopore Technologies, Oxford, UK). The amplified DNA samples were prepared according to the kit instructions and loaded onto the flow cell for sequencing following manufacturer’s instructions. Sequencing was carried out in Mk1B and GridION MK1 devices for 72 hours and the resulted fast5 files were basecalled using guppy v6.3.7 using the high accuracy model (dna_r9.4.1_450bps_hac).

Flye 2.9 (Kolmogorov et al. 2019) was used for assembly. For samples with >100X coverage, the “--asm-coverage 100” parameter was used to improve assembly and facilitate the assembler performance. We then used Nextpolish 1.4.1 (Hu et al. 2020) to increase the overall basecall quality of the genome and facilitate further quality control analysis such as Benchmarking Universal Single-Copy Orthologs (BUSCO) scores, and better gene annotation. Illumina sequencing was not used here because the objective of this research was to determine how MDA would affect long-read generation.

### Putative Concatemer Detection in Intact vs. Fragmented Amplified Samples

To examine the impact of fragmentation on MDA products, we treated DNA aliquots with NEBNext dsDNA Fragmentase (CAT# M0348S) at 37°C for 16 minutes, creating fragments between 500-1000 bp. This enzyme-based method induces DNA shearing, generating fragments of specified sizes in a time-dependent manner. The process provides random fragmentation similar to mechanical methods. Both fragmented and non-fragmented (high molecular weight DNA) samples were sequenced as described above.

The CADECT tool (https://github.com/rpbap/CADECT), was developed in-house and was used for the detection and removal of putative concatemeric chimeric sequences in the ONT amplified reads. CADECT splits all reads into separate files and performs sliding windows with a user-defined preferred size and gap between windows. For ONT amplified reads, a window size of >= 500 bp with no overlaps was used (*e.g*., -w 500 and -s 500). Reads generating less than one window (< 1 kb in size in the 500 bp window example) were skipped, and their IDs were stored in the short.txt output file. Fragment windows from reads with more than two windows were aligned using nucmer from mummer 4 (Marcais et al. 2018), and reads with overlaps were reported in the stats file, with their IDs stored in the concat_ID output file. Statistics including the total number of reads, number of putative concatemers, number of reads with no concatemer detection, and overlap frequency were recorded in the stats.txt output file. Fastq/Fasta files containing the characterized reads were generated for further analysis.

These methods were employed to investigate the benefits and limitations of multiple displacement amplification in whole-genome Oxford Nanopore Sequencing, focusing on low-concentration DNA samples.

### GC Bias Evaluation

To calculate GC bias in the sequencing depth of the amplified data, we compared the local %GC content using sliding windows of 1000 bp to the average coverage depth for each of these regions (https://github.com/DamienFr/GC_content_in_sliding_window). Depth windows were calculated using the R packages setDT and rollapply packages. R^2^ coefficients were calculated using the ordinary least squares regression method.

## DATA AVAILABILITY

The raw sequence data generated in this study have been submitted to the NCBI BioProject database (https://www.ncbi.nlm.nih.gov/bioproject/) under accession numbers PRJNA1063853 and PRJNA1022047. The assembled genomes in this project are preliminary drafts and are currently unavailable at this stage due to the scope of this project. They were generated solely using ONT reads and have not undergone polishing with Illumina short-read data. Additionally, they have not been checked for potentially contaminating “leftover” contigs. The raw data is accessible for reproduction purposes, and the final, polished, and decontaminated assemblies will be made available in subsequent publications.

## ACKNOWLEDGMENTS

This work was funded in part by National Institute for Allergy and Infectious Diseases, NIAID, R01 AI148667 to T.C.G. and J.C.K, NIH T32GM142623 to F.A.D. The funders played no role in the study design, data collection and analysis, decision to publish, or preparation of the manuscript.

## AUTHOR CONTRIBUTIONS

FAD, MSB, AD, HS, MQD, RPB performed the research. MSB, JCK, TCG, RPB conceived the research. FAD, MSB, AD, MQD and RPB analyzed the results. FAD, MSB, AD, RT, AK, CAA, JCK, TCG, RPB contributed to writing the manuscript. CAA, JCK, TCG and RPB obtained funding. All authors reviewed and approved of the manuscript.

## SUPPLEMENTAL FIGURES

**Figure S1.**
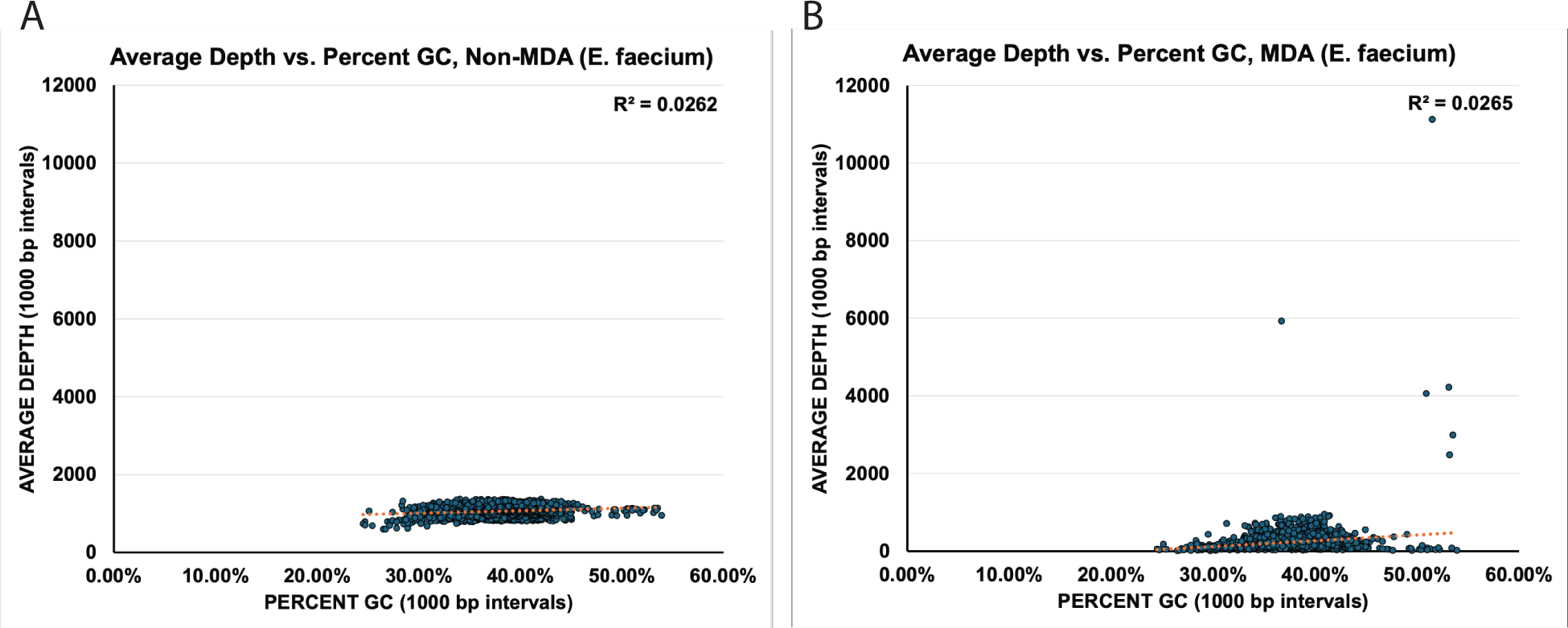
Square regression correlation coefficient between average depth and average %GC across 1000 base pair sliding window regions of the genomic assembly shows low correlation to %GC. A correlation analysis of (A) non-amplified and (B) amplified assemblies of *E. faecium*.

**Figure S2.**
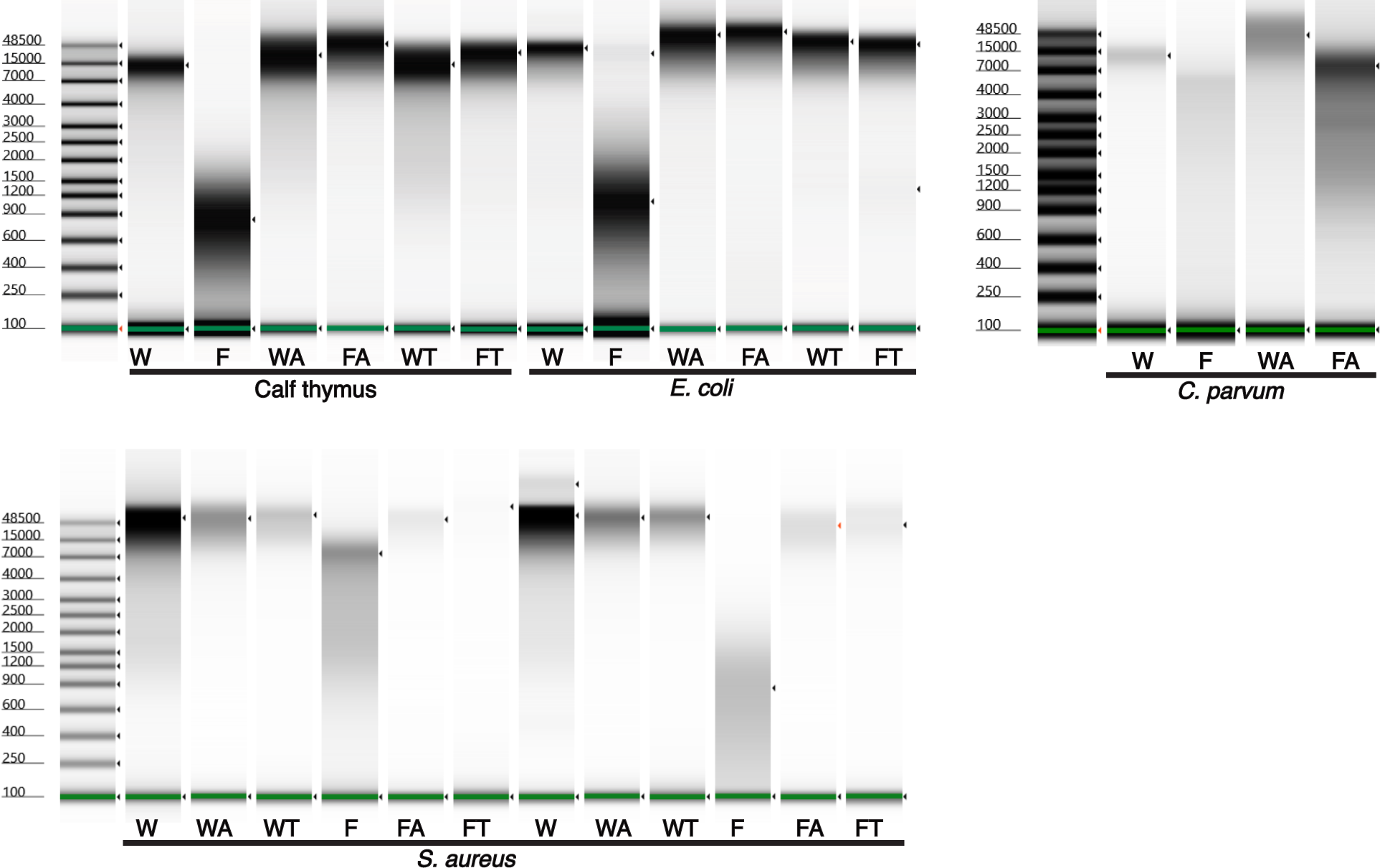
Uncropped DNA Fragment and read Size Range pre- and post-Multiple Displacement Amplification using Size-Controlled Fragmented DNA. Uncropped TapeStation results for different organism sets; (W = whole intact DNA; F = Fragmented DNA; WA and FA = after amplification; and WT and FT = after T7 debranching.

## SUPPLEMENTAL TABLES

**Table S1.**
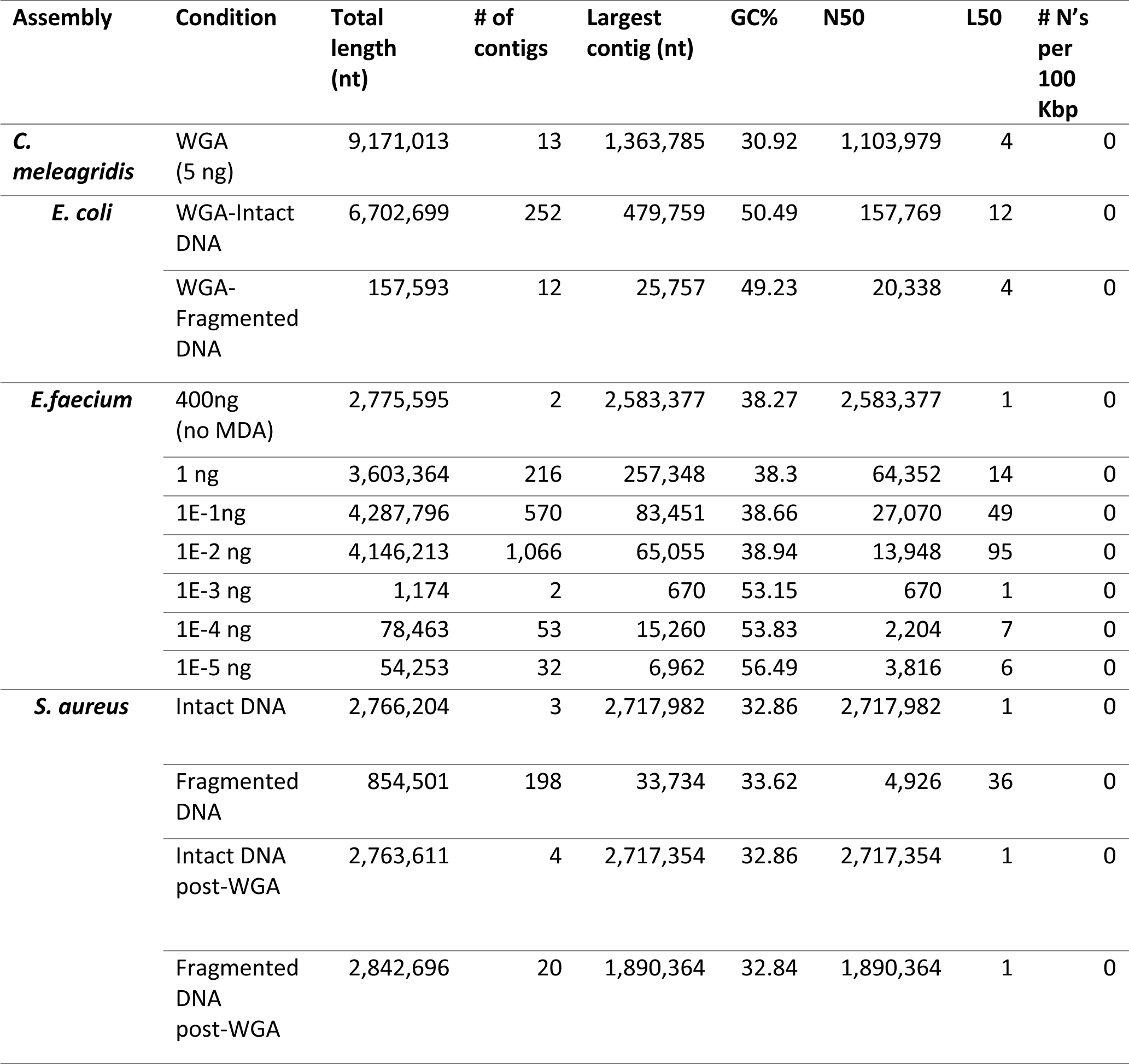
Whole genome assembly statistics for the data generated using MDA.

**Table S2.** Sequencing Statistics for *E. faecium\** WGS results at 2.5 ng and 0.2.5 ng starting DNA input.

| Starting total DNA | 2.5 ng | 0.25 ng |
| --- | --- | --- |
| Depth | 73× | 35× |
| Total length (bp) | 2,778,112 | 2,918,507 |
| Number of contigs | 3 | 35 |
| GC% | 38.2 | 38.28 |
| N50 | 2,589,111 | 233,792 |
| L50 | 1 | 4 |
| Number of N's per 100 Kbp | 0 | 0 |

\*the expected genome size for *E. faecium* species varies between 2.6 and 3.2 Mb

**Table S3.**
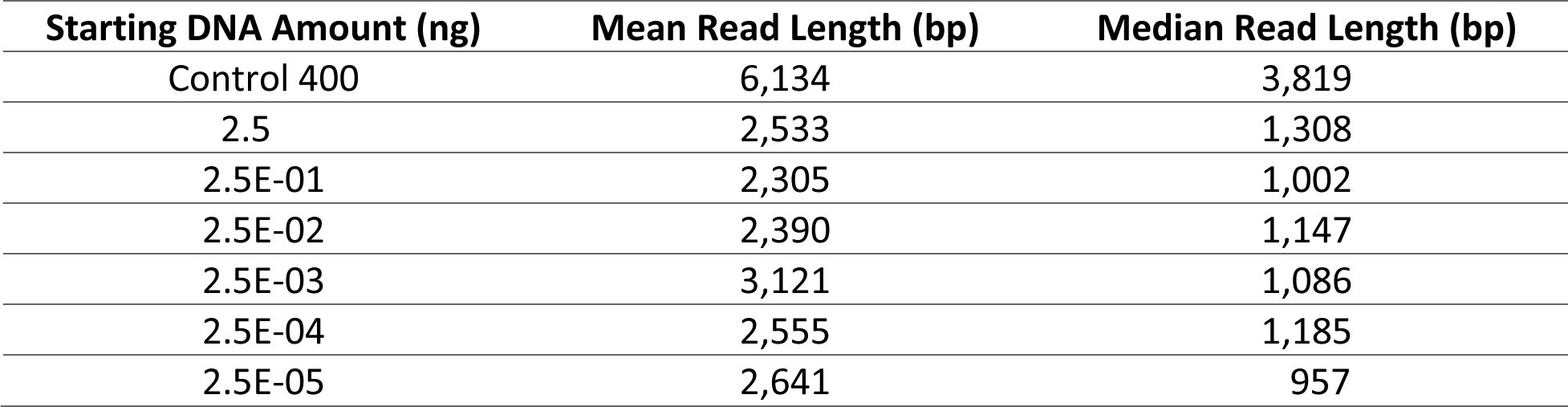
Read length distribution among the DNA dilutions for *E. faecium,* based on starting DNA concentration.

**Table S4.** Comparison between *S. aureus* ATCC-29213 genome sequences assembled before and after CADECT with assembly length, number of contigs, number of reads, and mean read length.

|  | Fragmented DNA with<br>MDA <u>before</u> CADECT | Fragmented DNA with<br>MDA <u>after</u> CADECT |
| --- | --- | --- |
| Assembly<br>length (bp) | 2,842,696 | 2,752,482 |
| Number of<br>contigs | 20 | 3 |
| Number of<br>reads | 324,197 | 292,789 |
| Mean read<br>length (bp) | 2,163 | 1,534 |
| Median read<br>length (bp) | 1,352 | 1,202 |

**Table S5.**
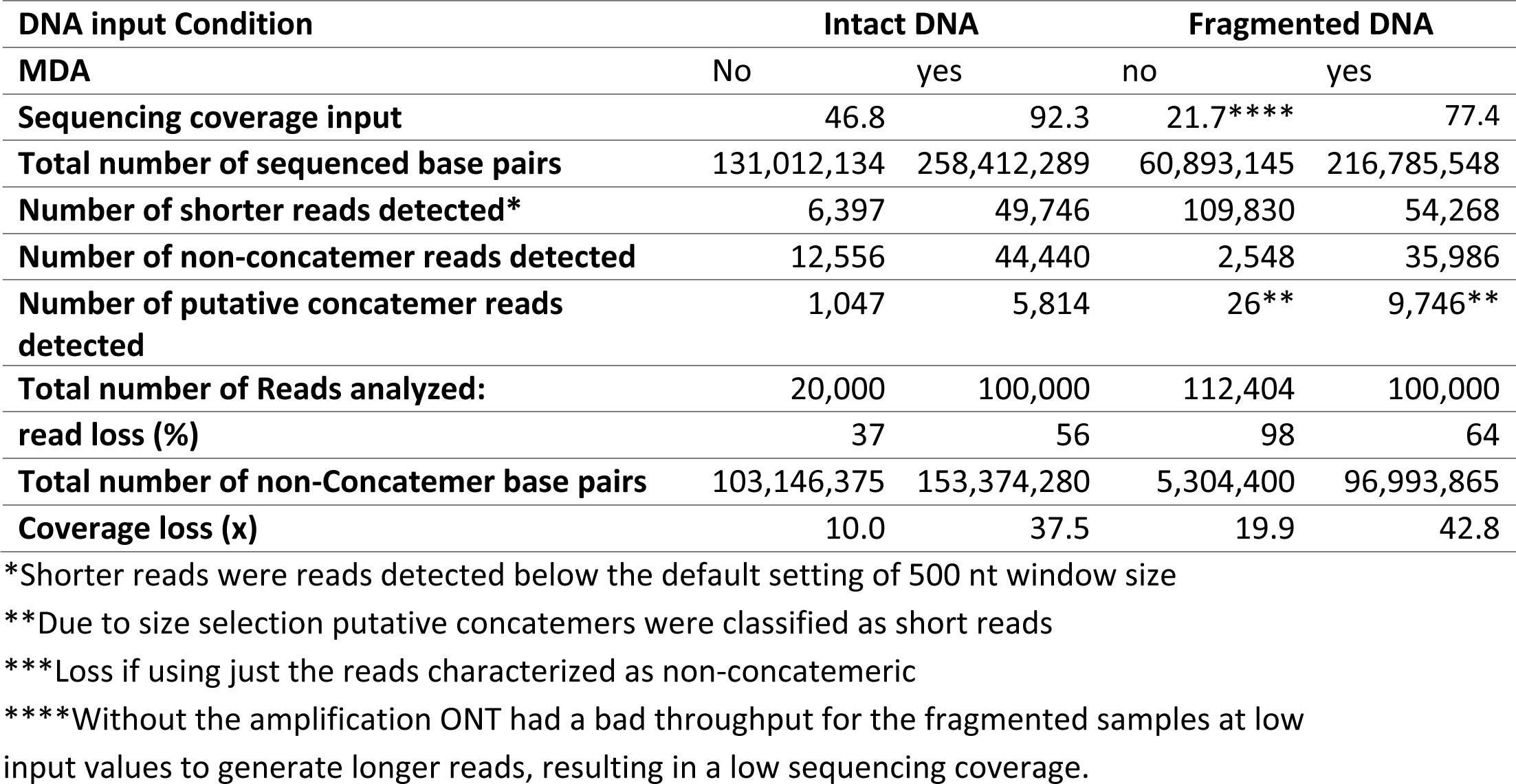
Comparison between *S. aureus* results from CADECT using low DNA input samples.

